# High-Capacity transcranial Direct Current Stimulation (HC-tDCS)

**DOI:** 10.1101/2025.06.11.659142

**Authors:** Kyle Donnery, Mohamad FallahRad, Niranjan Khadka, Matthew Saw, Benjamin Babaev, Santiago Osorio, Mojtaba Belali Koochesfahani, Rayyan Bhuiyan, Jana M. Elwassif, Marom Bikson

## Abstract

**Background:** Enhancing tDCS technology can support the delivery of higher current intensities, enabling broader dose-response studies in human trials.

**Methods:** High-Capacity tDCS (HC-tDCS) integrates novel electrodes and adaptive current/voltage controlled electronics. Multi-layer HC electrodes include polarity-specific redox layers and designed hydrogel interfaces, shaped for a bifrontotemporal montage. The stimulator design includes adaptive ramps with hybrid voltage-current control and low (7.5 V) compliance voltage. Scanning electron microscopy (SEM) and electrical impedance spectroscopy (EIS) were used to characterize electrode properties. Tolerability of HD-tDCS was tested for target currents 1-6 mA (in 1 mA increments) for 30 min on 5 healthy subjects, and compared with conventional tDCS using 2 mA F3-F4 sponge-electrodes. MRI-derived computational models predicted cortical electric fields. Tolerability was assessed according to the 100 mm visual analogue scale for pain (VASP-100), skin erythema assessment, thermal imaging, and adverse event questionnaires.

**Results:** The electrode design including high-roughness polarity-specific capacity, electrochemically supports high-charge direct current stimulation. In all subjects, HC-tDCS was well tolerated at all tested doses (1-6 mA) with minor transient adverse events and average VASP-100 less than 15. VASP-100 during sponge-electrode tDCS at 2 mA was comparable to 5 and 6 mA HC-tDCS. HC-tDCS operates at significantly lower voltage than sponge-tDCS, impacting tolerability and efficiency. Modeling predicts peak frontal electric fields of 0.65-1.08 V/m for 2 mA HC-tDCS and 1.95-3.25 V/m for 6 mA HC-tDCS, compared to 0.49-0.95 V/m for 2 mA sponge-tDCS.

**Conclusions:** HC-tDCS allows increased cortical stimulation; at 6 mA achieving double the 1 V/m electric field threshold in all subjects. Enabled by pre-stimulation procedures, specialized electrodes, and adaptive low-voltage stimulators, HC-tDCS is well tolerated at intensities up to at least 6 mA.

## Introduction

The majority of transcranial direct current stimulation (tDCS) trials used an intensity of 2 mA (1,2). Relatively few studies explored higher intensities, ranging between 2.5 and 4 mA per electrode (3–8) or employed multi-electrode (High-Definition) montages to increase total current delivery.

Increasing intensity increases brain electric fields linearly (9) and enhances cellular responses in animal models (10–13). tDCS clinical response has been correlated with higher individual brain electric fields (14–16).

As tDCS with a few mA is well below accepted safety limits (1), the reluctance to explore higher current derives from convention and concerns around adverse events at the skin (tolerability). Improved electrode designs mitigate skin effects including High-Definition (HD) electrodes (17), dry electrodes (18,19), as well as sponge electrode salinity (20–21) and rivets (22).

This study introduces High-Capacity tDCS (HC-tDCS) electrodes and stimulators, designed to deliver up to 6 mA in a bifrontotemporal montage without compromising tolerability compared to 2 mA sponge-electrode tDCS. The 6 mA HC-tDCS electric fields exceed 2 V/m. Designed features include polarity-specific roughend electrode redox-layers, low-voltage (7.5 V) adaptive intensity control, and pre-stimulation procedures including skin-conditioning wipe.

## Methods

### Electrodes and Device Specifications

Stimulation was delivered using a custom stimulator designed with hybrid current/voltage control circuitry. The device was programmed to execute an adaptive, voltage-controlled ramp over 60 s with a current limit set to the target current (1-6 mA). A design specification was achieved >90% of the target current in <90 s (including 60 s of ramp) across all subjects. Following the adaptive ramp period, constant-current stimulation was maintained for 30 min, followed by a linear voltage-controlled ramp down. The device, electrode design, and preparation techniques were centered on minimizing the stimulator compliance voltage needed to achieve dose specification, motivated by 1) reducing voltage changes at the redox electrolyte (electrochemical overpotential), the skin-electrolyte interface, and across skin superficial layers; 2) optimized hardware requirements and power (23).

For HC-tDCS, the stimulator was configured to deliver target currents ranging from 1 – 6 mA with a compliance voltage of 7.5 V. For sponge tDCS, the stimulator was configured to deliver 2 mA with a compliance voltage of 20 V (24), which is still below the compliance voltage of conventional tDCS devices.

### Electrode Fabrication

HC-tDCS electrodes (Fig. 1a) were fabricated using an additive layered manufacturing process (Fig. 1b), integrating five functional layers into a compact (∼0.8 mm thick), high-performance assembly. The electrode topology was specificallydesigned for a bifrontotemporal stimulation montage, targeting the bilateral (pre)frontal cortex. The electrode shape is a novel falcate triangular geometric profile, which maximizes the surface contact area across the curvature of the frontotemporal scalp.

**Figure 1.**
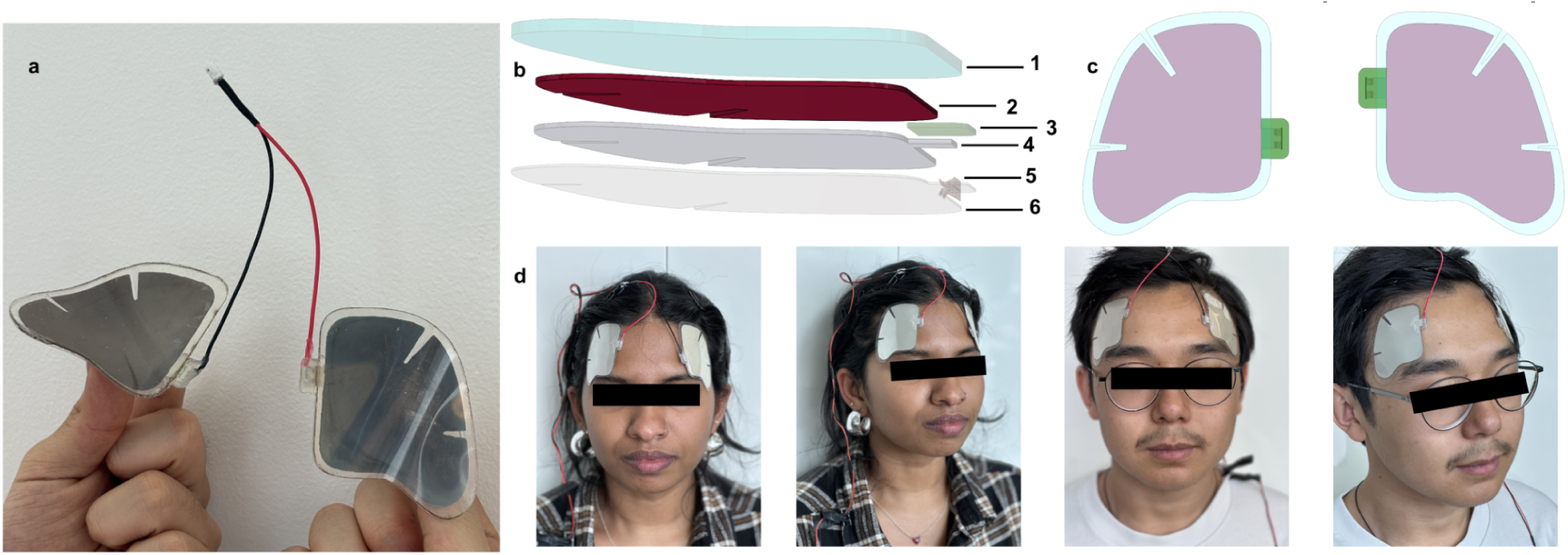
| Design and application of High-Capacity tDCS (HC-tDCS) electrodes. **a**) Photograph of the HC-tDCS electrode pair, illustrating the assembled electrodes with attached leads. **b**) Schematic cross-section of the multilayer electrode design (from top to bottom): (1) adhesive hydrogel layer, (2) electrode (redox) layer, (3) dielectric layer, (4) conductive layer, (5) metal connector for the lead wires, and (6) substrate layer. For visualization purposes, the thickness of each layer (excluding layer 5) has been increased by 10x. **c**) Plan view rendering of the electrodes highlighting placement of the strain-relief slits. **d**) Representative images of participants with HC-tDCS electrodes positioning for the bifrontotemporal montage, showing both frontal and lateral views.

Conformability, especially along the brow ridge and hairline, was enhanced with two strain relieving slits in the electrode. Note that these slits are present in all layers except the hydrogel. The electrode contact surface was 25 cm², matching 5 cm x 5 cm sponge-electrode pads. The layered electrode assembly consists of the following components: (1) an outer ion-conductive hydrogel layer, which functions as the skin interface and adhesion mechanism, eliminating the need for a headgear securing mechanism; (2) a non-conductive acrylic polymer insulating layer applied to shield non-stimulating regions, preventing exposure of metal parts; (3) an anode– or cathode-specific redox layer—zinc electroplated onto the anode or AgCl formed through controlled corrosion on the cathode—to enhance the redox capacity at the electrode-hydrogel interface; (4) a conductive silver layer, applied by printing and cured to ensure efficient transmission of electrical current from the wired connector; and (5) a conductive clip (S0971-46R, DigiKey, Thief River Falls, United States), functioning as an conductive connection for the leads; (6) a 0.1 mm polyethylene terephthalate (PET) substrate providing mechanical integrity for the electrode. The hydrogel layer has a volume resistivity of ≤ 600 Ω·cm, a controlled uniform thickness of 0.6 mm (±0.1 mm) and a mildly acidic pH of 3.5 (±0.5). The hydrogel’s water activity (∼0.66) preserves hydration and supports long-term ionic conductivity, minimizing impedance drift over the course of stimulation sessions. The hydrogel is made using non-cytotoxic, non-irritating, and non-sensitizing materials.

### Scanning Electron Microscopy (SEM)

For characterization of the electrode surface, electrode samples without the hydrogel layer were prepared on conductive platforms. SEM images were acquired using a Zeiss Supra 55 field emission scanning electron microscope (ZEISS, Oberkochen, Germany) operated at 5 kV accelerating voltage and a 4 mm working distance, using a secondary electron detector.

### Electrode Impedance Spectroscopy (EIS) and Overpotential

EIS measurements were performed using a frequency response analyzer (FRA51615, NF Corporation, Yokohama, Japan) over a frequency range of 10⁻² Hz to 10² Hz, with 30 logarithmically spaced frequency steps. A 10 mV AC excitation waveform was applied, and the DC bias was dynamically adjusted to compensate for shifts in the open-circuit potential of the electrode system before and after stimulation. The experimental setup consisted of an anode and cathode connected via a hydrogel interface, forming a closed two-electrode configuration. All measurements were conducted in a controlled environment to ensure measurement stability. Baseline impedance spectra were collected prior to stimulation, after which a 10 mA constant-current discharge was applied for 60 min. The system was then allowed to equilibrate for 15 min to enable electrochemical stabilization before post-discharge EIS measurements were acquired under identical conditions.

Overpotential measurements were conducted using the same electrode configuration as employed in the EIS protocol, with the anode and cathode directly interfaced via a hydrogel layer. Constant-current discharges were performed at 1 to 10 mA in 1 mA increments, each for a duration of 1 hr. Voltage and current were recorded using a source meter (2450 SMU, Keithley Instruments, Beaverton, United States) operating in 2-wire mode, with data sampled every 0.5 s throughout the discharge period. To facilitate comparison across conditions, all traces were baseline-shifted such that overpotential at the start of each discharge was normalized to 0 V.

### Computational modeling and Solution Method

Eight high-resolution (1 mm^3^) healthy human subjects’ magnetic resonance imaging (MRI) scans were segmented into six masks namely scalp, skull, cerebrospinal fluid (CSF), grey matter, white matter, and air using Simpleware (Synopsys Inc, CA, USA) to develop a finite element method (FEM) model using series of automatic and manual morphological tools. Computer aided design (CAD) models of 35 cm^2^ sponge pad and 25 mm^2^ hydrogel electrode were first designed in Solidworks (Dassault Systems Americas Corp., MA, USA) and later imported and positioned over F3-F4 region (sponge pad) and bifrontotemporal region (hydrogel electrode) in Simpleware. Using built-in voxel-based meshing algorithms in Simpleware, we generated an adaptive tetrahedral mesh. The mesh density was refined until additional model refinement produced less than 1% difference in electric field (electric field) at the grey matter. The resulting volumetric mesh was then imported to COMSOL Multiphysics 6.1 (COMSOL Inc., MA, USA) to computationally solve the electric fields distribution across models. The segmented masks were assigned an isotropic electrical conductivity (S/m): scalp: 0.20 S/m (combined skin and fat conductivity); skull: 0.01 S/m; CSF: 0.85 S/m; grey matter: 0.265 S/m; white matter: 0.126 S/m; air: 1×10^-6^ S/m (25–29). The corresponding conductivity of sponge-electrodes and hydrogels were assigned as 1.4 S/m (17) and 0.166 S/m, respectively.

tDCS was simulated by solving the Laplace equation (∇ (σ∇V) = 0 where ‘V’ is potential, ‘σ’ is conductivity) under quasi-static assumption (26,30) to predict electric field magnitude at the cortex. The boundary conditions were applied as normal current density (inward current flow: J_norm_) at the top exposed surface of anode electrode (2 mA for F3-F4 montage with sponge-electrode; 1, 2, 3 4 5 6 mA for bifrontotemporal montage with hydrogel-based electrode) while the top surface of cathode electrode was grounded (0 mA) to represent different montages for all head models (or alternatively 1 mA was applied and the electric field scaled accordingly). The interior boundaries were assigned continuity (n⋅J = 0; n: surface normal; J = current density) and all other external boundaries were electrically insulated (n⋅J = 0). The final FEM models comprised >30 M tetrahedral elements. The relative tolerance was set to 1×10^-6^ to improve the solution accuracy. Peak electric field magnitude was predicted at the cortex for both montages using different electrodes and at various intensities. Peak electric field magnitude represents the 99^th^ percentile of electric field magnitude (31) produced in a 2×2×1 mm^3^ region of interest at the cortex local electric field maximum.

### Subject Testing Study Protocol

The study was conducted in accordance with protocols and procedures approved by the Institutional Review Board of the City College of New York. All volunteer participants provided written informed consent to participate in the study. Participants were excluded if they presented with or reported eczema, severe rashes, blisters, open wounds, burns including sunburns, cuts or irritation, or other skin defects which compromise the integrity of the skin at or near stimulation sites. Five healthy male subjects between the ages of 18 yrs and 49 yrs (M = 34 yrs, SD = ±9.3) were recruited and financially compensated for their participation.

Participants testing was conducted using a single-blind, randomized crossover design. Each participant underwent a total of seven stimulation sessions including HC-tDCS at target intensities ranging from 1 – 6 mA, with 1 mA increments, for 30 min (excluding adaptive ramp time) with the order of dose administration counterbalanced across subjects using *urn randomization* (32). Following the 6 HC-tDCS stimulation sessions, participants completed one session using 25 cm^2^ conventional sponge-electrodes using a bifrontal (F3-F4) headgear (SNAPstrap, Soterix Medical, New York, NY, USA) at 2 mA for 30 min (excluding ramp time). Each pair of sponge-tDCS and HC-tDCS electrodes (anode and cathode) was used once (i.e. single use). All sessions were right-anode polarity. Sessions were limited to one per day.

Prior to HC-tDCS stimulation, the stimulation site was gently wiped with a custom skin wipe made from nonwoven felt saturated with surfactant, alcohol, water and electrolyte. The wipe is designed to reduce skin inhomogeneity and impedance.

Pain ratings were collected using a visual analog scale (VAS-100) at baseline, after the 1 minute ramp, every 2 minutes during the 30-min stimulation period, and after stimulation concluded.

Subjects were allowed to terminate the session at any time. If subjects reported a VASP-100 scores ≥ 60 study or VASP-100 ≥ 50 for two consecutive blocks (2 min) the session would be aborted. These conditions did not occur in any stimulation sessions.

### Lexical Decision Task Distractor

During each stimulation session, participants completed a lexical decision task (LDT) as a distractor. The LDT was presented on a screen for 30-min and was composed of 15 two-min blocks. Each block contained randomized presentations of words and pseudowords, with response time and accuracy recorded. Task presentation and behavioral data were managed and analyzed using custom Python scripts, with particular attention to dose-dependent effects on cognitive performance.

### Outcome Measures

Prior to and following each session, participants completed the Positive and Negative Affect Schedule (PANAS) plus 3 Visual Analogue Mood scales (Happiness, Sadness, and Anxiety) and a standard tDCS Adverse Effects Questionnaire (33) to measure mood changes and tolerability. Thermal (Kaiweets KTI-W01 Therma) and digital images of the forehead (including region where electrodes contact skin), were captured prior to and after stimulation sessions under controlled lighting conditions. Skin erythema was characterized using the Draize erythema scoring system scale (34).

### Statistical Analysis

To model both the dose effects observed among the HC-tDCS trials and the differences between electrode tolerability and performance when controlled for current (comparing 2 mA conventional and 2 mA HC-tDCS), we used linear mixed-effects models, fitted with the “*lme4*” package in R (Posit, Boston, United States). Models used for the analysis of the effect of current (1 – 6 mA) on HC-tDCS outcomes utilized fixed effects for current (continuous), time (continuous), and their interaction (current times time). Whereas the models used for the analysis of differences between sponge tDCS and HC-tDCS utilized type (quantized), time (continuous), and their interaction (type times time) as fixed effects. When comparing sponge electrodes to HC-tDCS controlled for current, we used sponge electrodes as the reference for the model. For comparison of skin erythema and temperature, the models used current (continuous), time(continuous), and side (quantized) to account for variations in behavior under the anode and cathode. The Satterthwaite approximation estimated the degrees of freedom for the fixed effects, which provides more accurate significance testing by accounting for variability in smaller samples, thus minimizing the risk of Type I errors. All statistical analyses were performed in R. The significance level (α) was set at 0.05 and 95% confidence intervals were calculated for each of the fixed effects.

## Results

### HC Electrode Redox Layer SEM

SEM analysis of the anode, AgCl, (Fig. 2a-c) and cathode, Zn, (Fig. 2d-f) HC electrode redox layers demonstrated increased surface roughness (functional area). 2 cm x 3 cm samples were prepared of the metal and redox layers (Fig. 2a.1, 2b.1) appearing uniform and planer. High resolution SEM images show a (Fig. 2a.2,3, Fig. 2b.2,3) rough three-dimensional surface topology, which increases the real surface area available for redox functions (Fig. 2a.2,3, Fig. 2b.2,3).

**Figure 2.**
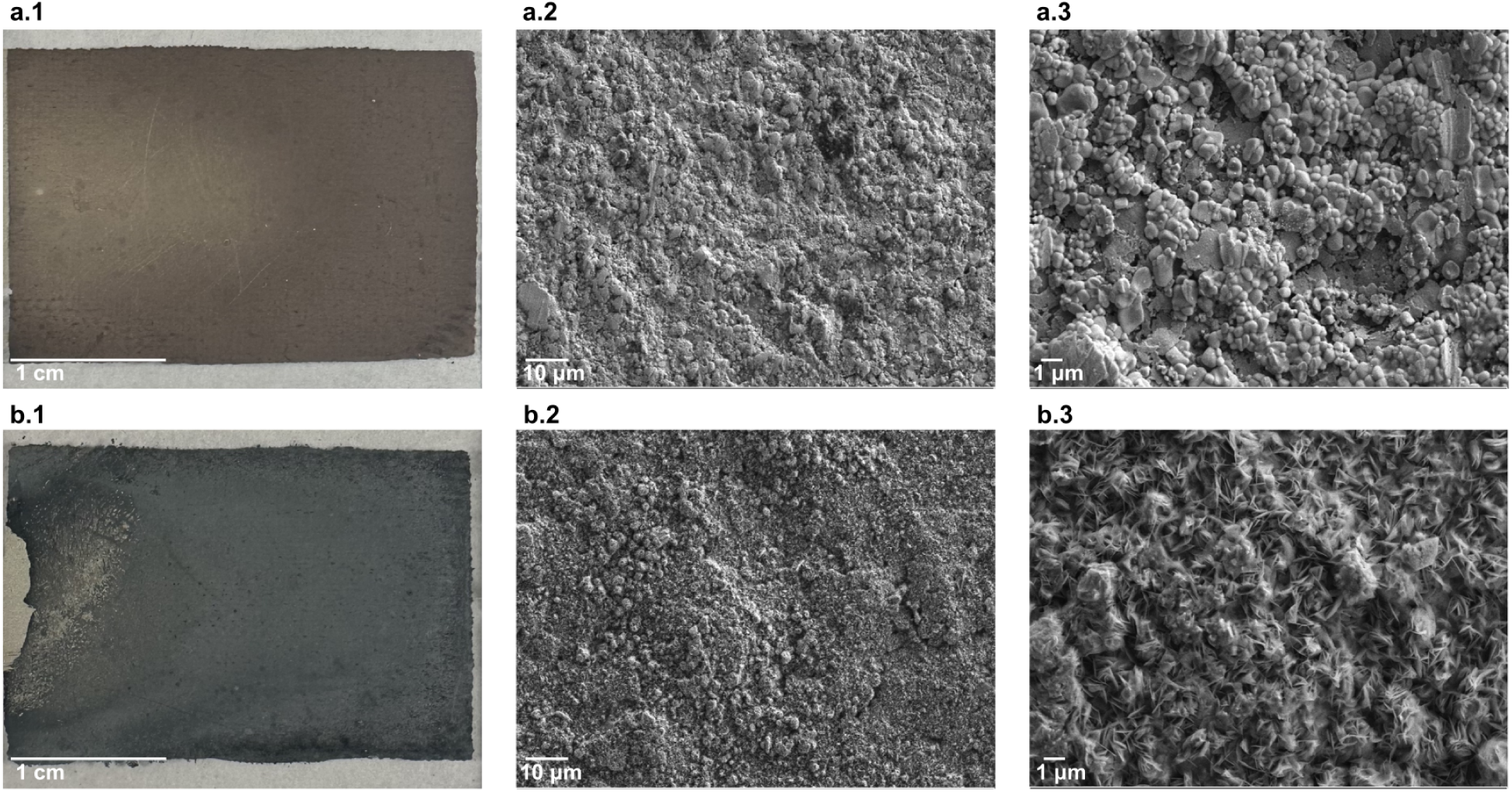
| Surface morphology of High-Capacity (HC) electrode redox layers. Imaging of electrode redox surface (top: anode, Zn;panel bottom cathode, AgCl) from optimal macroscale (1 cm scale bar) (a.1, b.1) to SEM microscale (a.2 and b.2 (10 μm scale bar) and a.3 and b.3 (1 μm scale bar)) of an unstimulated 3 cm × 2 cm electrode samples.

**Figure 3.**
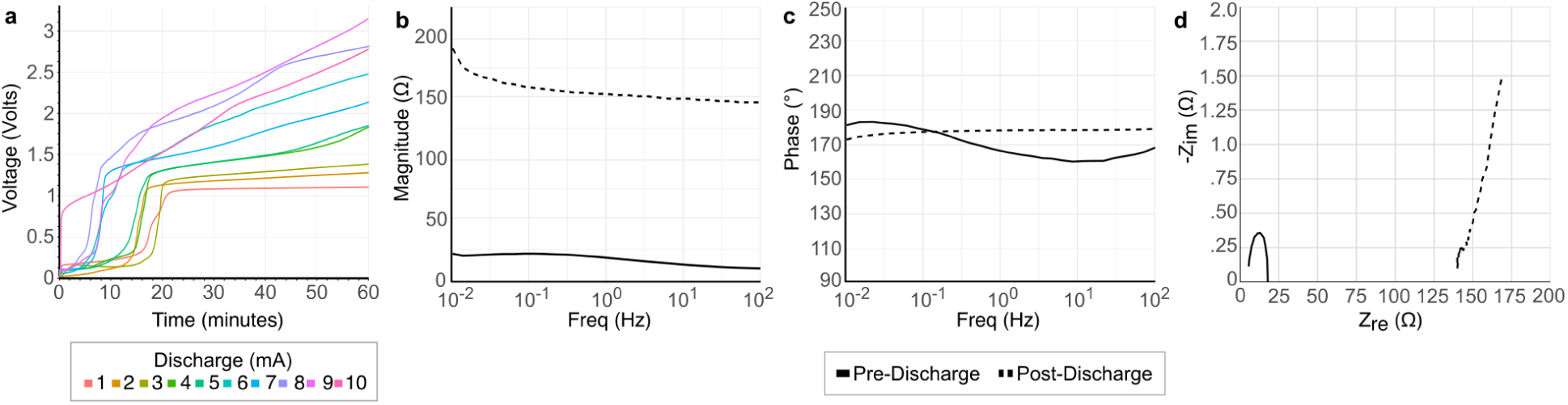
| HC electrode impedance during high current discharge. (a) Overpotential during constant current discharge at 1–10 mA. (a-c) Impedance pre– (solid lines) and post– (dashed line) 10 mA 60 min stimulation, showing magnitude (b), phase (c), and Nyuist plots (real: Z_re_ and imaginary: –Z_im_ components)

### HD Electrode EIS and Overpotential

HC electrode electrical performance was accessed by stimulating two HC electrodes connected at their hydrogels. In the first test, 10 mA 60 min direct current was applied (in excess of HC-tDCS specification); impedance was measured using EIS before and after stimulation. Electrode impedance magnitude increased after stimulation, consistent with redox layer depletion, with impedance phase largely unchanged (Fig b-d).

In a separate second test, two HC electrodes were stimulated 1 to 10 mA (in 1 mA increments) for 60 minutes and overpotentials were recorded. Potential increases depending on current and time. There is generally of low (<0.2 V) overpotential for an initial phase, followed by an voltage inflection to ∼1V (reflecting exhaustion of the first electrochemical reaction capacity of the redox electrode layer) for a second phase (a second electrochemical process), followed their phase by a linear increase in voltage (consistent with redox breakdown and capacitive charging). For HC-tDCS doses within specification (up to 6 mA, 30 min) the third phase did not occur.

### Brain Current Flow Computational Modeling

Cortical electric field magnitudes were predicted in eight MRI-derived head models under the conventional F3-F4 tDCS montage at 2 mA and the bifrontotemporal HD-tDCS montage at 1-6 mA. One an exemplary head, the model anatomy and cortical electric field are shown for 2 mA F3-F4 tDCS (Fig. a.1,2), 2 mA HC-tDCS (Fig. b.1,2) and 6 mA HC-tDCS (Fig. b.3). As expected (34), peak electric fields varied across subjects reflecting individual anatomy (Fig. 4c). The peak electric field ranges were: 2 mA F3-F4 tDCS 0.50 – 0.85 V/m; 1 mA HC-tDCS 0.33 – 0.50 V/m; 2 mA HC-tDCS 0.65 –0.99 V/m; 3 mA HC-tDCS 0.92 – 1.49 V/m; 4 mA HC-tDCS 1.32 – 1.98 V/m; 5 mA HC-tDCS 1.63 – 2.48 V/m; 6 mA HC-tDCS 1.96 – 2.99 V/m.

For the same applied current (2 mA), bifrontotemporal HC-tDCS results in marginally higher electric fields than F3-F4 tDCS, reflecting the electrode placement. For each subject, electric fields scale linearly with current applied. At 3 mA HC-tDCS, the subject with the least electric field exceeds the subject with the most electric fields for 2 mA F3-F4 tDCS. At 4 mA HC-tDCS, every subject has electric fields >1 V/m. At 6 mA HC-tDCS, the subject with the least electric field has double the electric field of the subject with the highest electric field for 2 mA F3-F4 tDCS.

### Human Trial: Pain, Voltage and Impedance

HC-tDCS at all target currents (1-6 mA) was well tolerated by participants with average VAS-P100 score during stimulation less than 15 (Fig. 5a.3). There was a significant main effect of current-intensity (β = 2.71, SE = 0.30, t(488) = 8.89, p < 0.001) and time (β = –0.21, SE = 0.06, t(488) = –3.32, p = 0.001) on pain ratings, and a significant current-intensity and time interaction (β = –0.05, SE = 0.02, t(488) = –3.29, p = 0.001). Higher currents were associated with moderately increased reported pain, while pain ratings decreased over time (Fig. 5a.1). At the first 2 minutes after the ramp, when pain sensations are expected to be higher, the median reported pain was less than 20, even at the 6 mA intensity (Fig. 5a.2).

**Figure 4.**
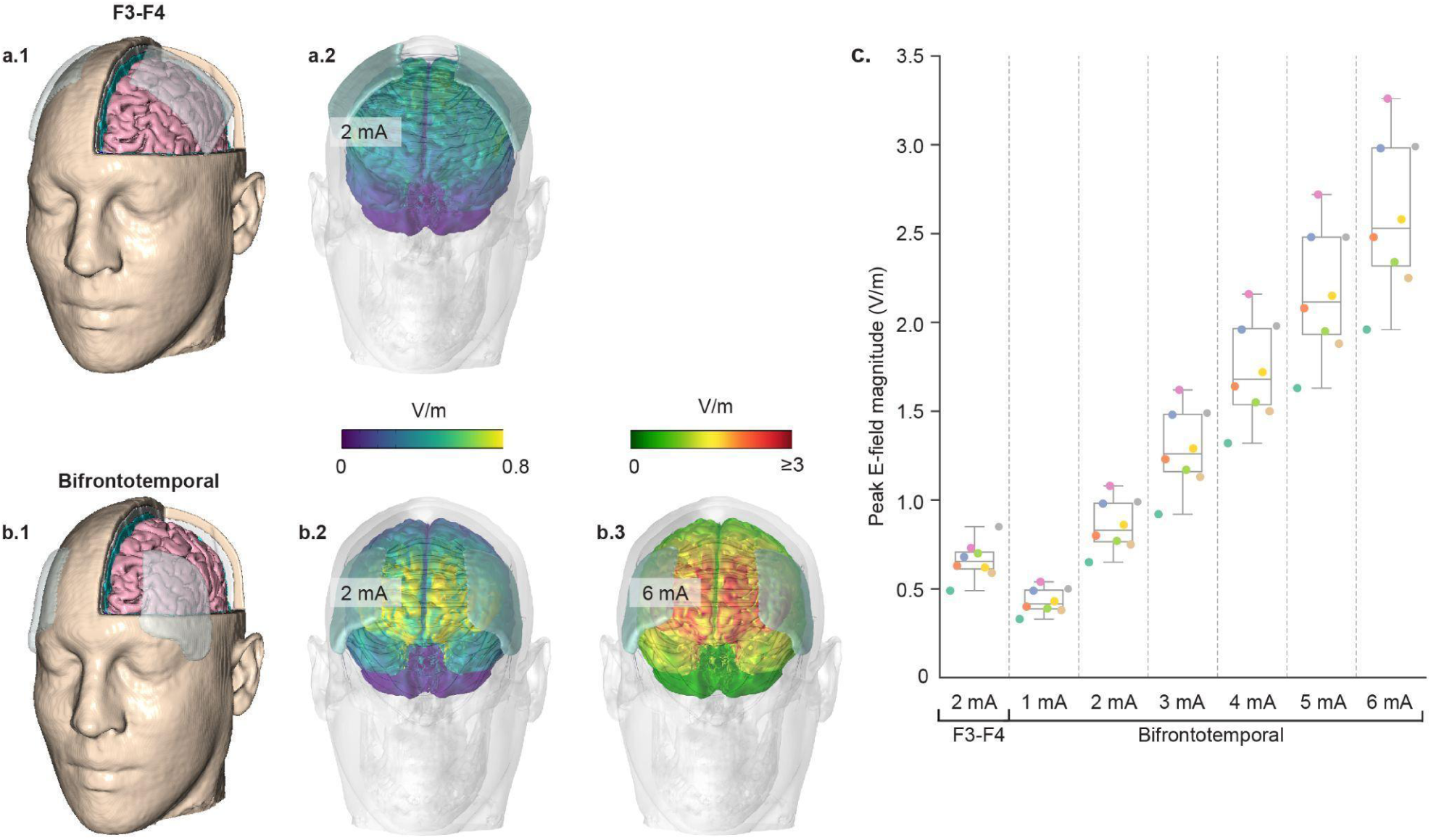
| Predicted brain electric fields magnitude during F3-F4 tDCS at 2 mA and bifrontotemporal HC-DCS at 1-6 mA across eight subjects. (**a.1, b.1**) 3D cut-view of exemplary subject anatomy with the sponge electrodes positioned at F3-F4 and HC electrodes positioned at bifrontotemporal positions. For an exemplary subject, cortical electric fields at 2 mA conventional tDCS (**a.2**), 2 mA HC-tDCS **(b.2**), and 6 mA HD-tDCS (**b.3**). electric field shown in false color; note distinct scale for 6 mA HC-tDCS. (**c**) Distribution of peak electric fields across subjects and montages/current intensities. Box plots with each subject color coded.

**Figure 5.**
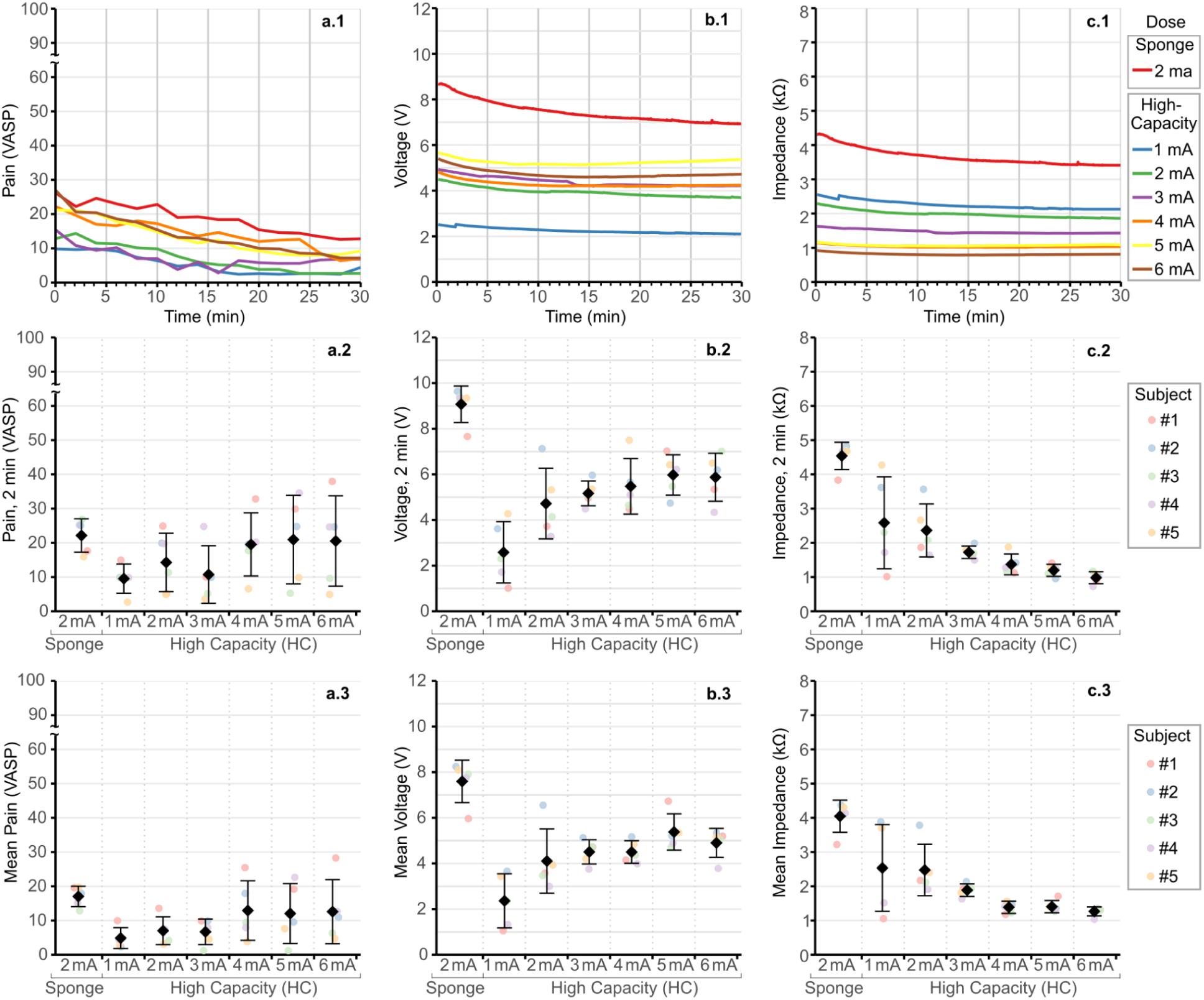
| Subject reported pain (VAS-P100), voltage, and impedance during HC-tDCS (1-6 mA) and conventional tDCS (2 mA). (a.1). Average VAS-P100 for each dose (line color) over 30 min of stimulation (**a.2**) VAS-P100 2 min after start of stimulation (after end of ramp up) for each subject (point color). (**a.3**) Average VAS-P100 during session for each subject (point color). **(b.1**) Stimulator output voltage for each dose (line color) over 30 min of stimulation (**b.2**) Stimulator output voltage 2 min after start of stimulation (after end of ramp up) for each subject (point color). (**b.3**) Average stimulator output voltage during session for each subject (point color). **(c.1**) Impedance for each dose (line color) over 30 min of stimulation **(c.2**) Impedance 2 min after start of stimulation (after end of ramp up) for each subject (point color). **(c.2**) Average impedance during session for each subject (point color).

**Figure 6.**
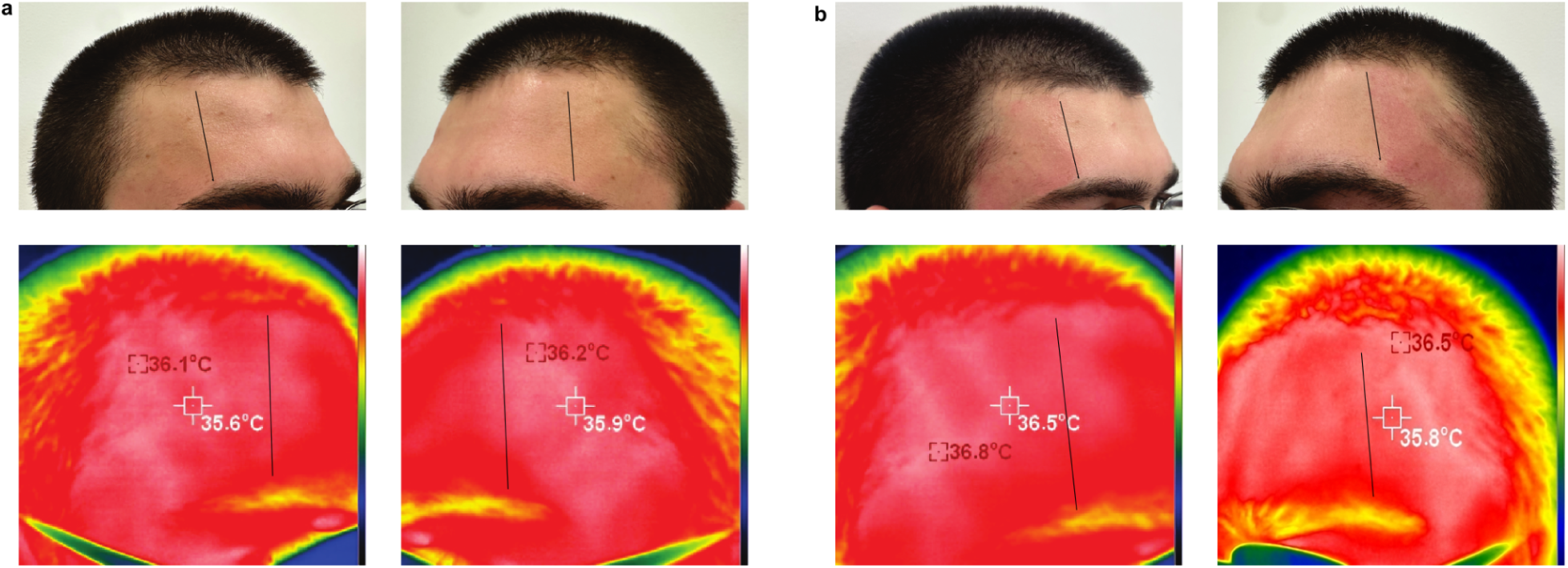
| Optical and thermal photography of stimulation site before (a) and after (b) stimulation. Solid black lines are used to represent the medial plane of the HC-tDCS electrode when placed on the forehead for stimulation.

Comparing HC-tDCS at 2 mA to conventional sponge-electrode tDCS at 2 mA, there was a significant main effect of condition in pain score (β = –12.01, SE = 1.44, t(168) = –8.35, p < 0.001); at 2 mA average pain scores for HC-tDCS were 12 points lower than conventional tDCS. A significant main effect of time (β = –0.45, SE = 0.06, t(168) = –7.88, p < 0.001), indicated that pain ratings decreased over time across both conditions. There was no significant current-by-time interaction in this comparison. There was a significant overlap between the VASP-100 scores reported by participants after ramp up during the 2 mA sponge tDCS and 2-6 mA HC-tDCS (Fig. 5a.2).

As target current intensity of HC-tDCS sessions increased, there was a moderate increase in stimulator output voltage required to maintain the target current (Fig. 5b.1). There was a significant main effect of current intensity for voltage (β = 0.46, SE = 0.003, t = 137.0, p < 0.0001). Time was also a significant predictor (β = –0.00038, SE = 0.000011, t = –34.32, p < 0.0001), with voltage decreasing over time (Fig 5 b.1). The current and time interaction was significant (β = 1.05 × 10⁻⁵, SE = 2.83 × 10⁻⁶, t = 3.71, p = 0.0002), suggesting less voltage reduction over time for higher currents. When directly comparing the sponge-electrode tDCS and HC-tDCS, voltages were significantly higher during sponge-electrode stimulation (β = –4.43, SE = 0.02, t = –238.32, p < 0.0001, reference: sponge).

HC-tDCS required less voltage at any current (1-6 mA) than sponge-electrode tDCS at 2 mA. The average voltage to maintain 6 mA HC-tDCS was ∼1.6X less than the average voltage required to maintain 2 mA sponge-electrode tDCS (Fig. 5.b3). In the linear mixed effect model, time remained a significant negative predictor (β = –0.00145, SE = 0.000013, t = –113.85, p < 0.0001), indicating a decrease in voltage over time for both conditions. A significant electrode-type and time interaction was observed (β = 0.00107, SE = 0.000017, t = 64.38, p < 0.0001), indicating that the rate of voltage decrease over time differed for HC-tDCS and sponge-electrode tDCS.

For impedance across HC-tDCS session, there were significant main effects of target current (β = –336.7, SE = 8.5, p < 0.0001) and time (β = –65.2, SE = 1.8, p < 0.0001), as well as a significant current-by-time interaction (β = 3.44, SE = 0.46, p < 0.0001). Impedance was also significantly lower in HC-tDCS as compared to sponge tDCS (β = –1264.7, SE = 51.8, p < 0.0001). Both time (β = –72.5, SE = 2.1, p < 0.0001) and the current-by-time interaction (β = –9.7, SE = 2.9, p = 0.0007) were significant.

### Human Trial: Adverse Events and Skin Response

All sessions of HC-tDCS were well tolerated with transient and mild sensations. Table 1 summarized the intensity and attribution scores for adverse events before and after HC-tDCS at 1-6 mA current intensities as well as conventional sponge-electrode tDCS at 2 mA. Baseline symptom intensities were uniformly low with mean scores near 1.0, indicating “absent” symptoms. Following stimulation, mean intensity and relation ratings did not change or increased incrementally.

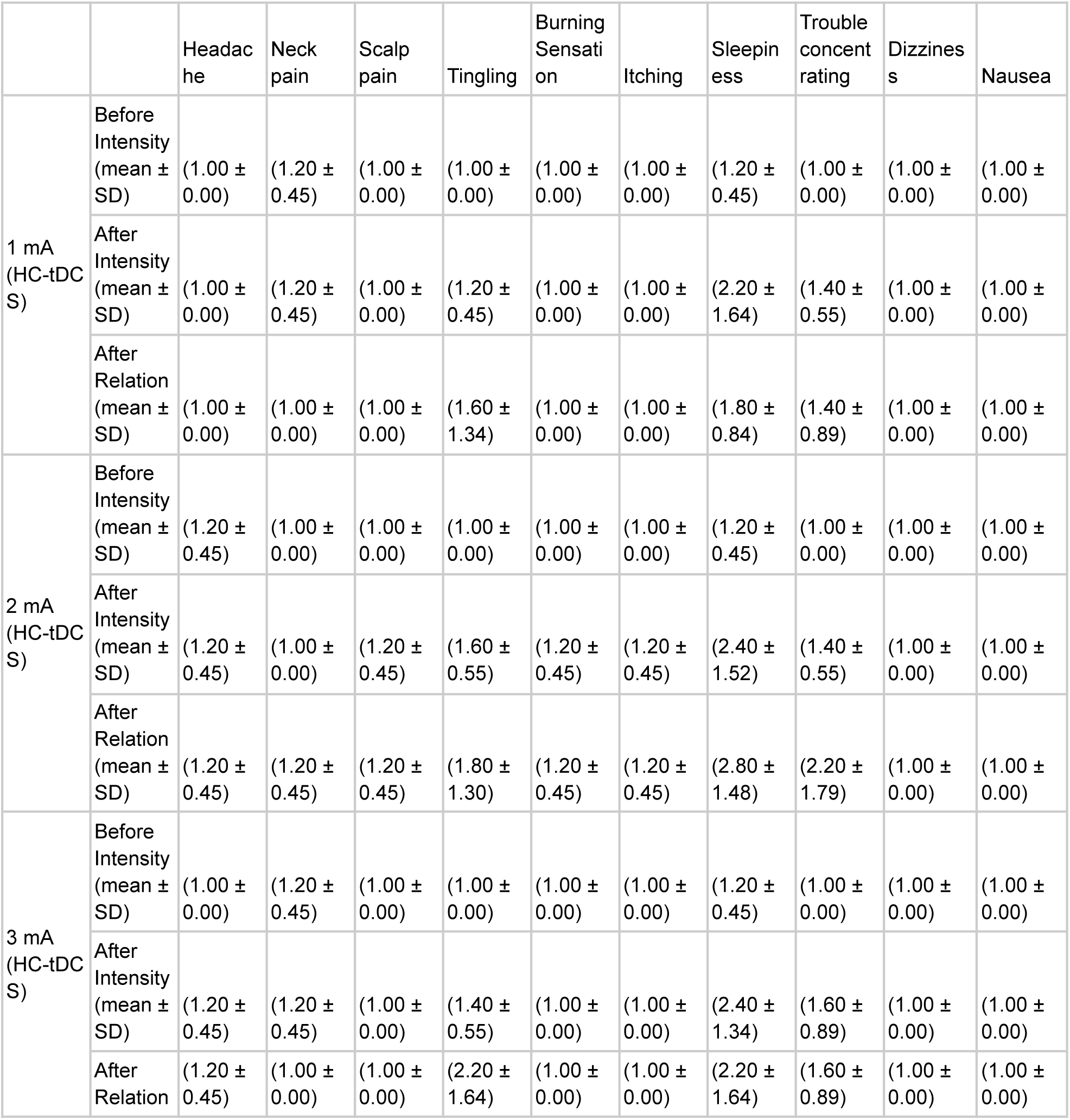
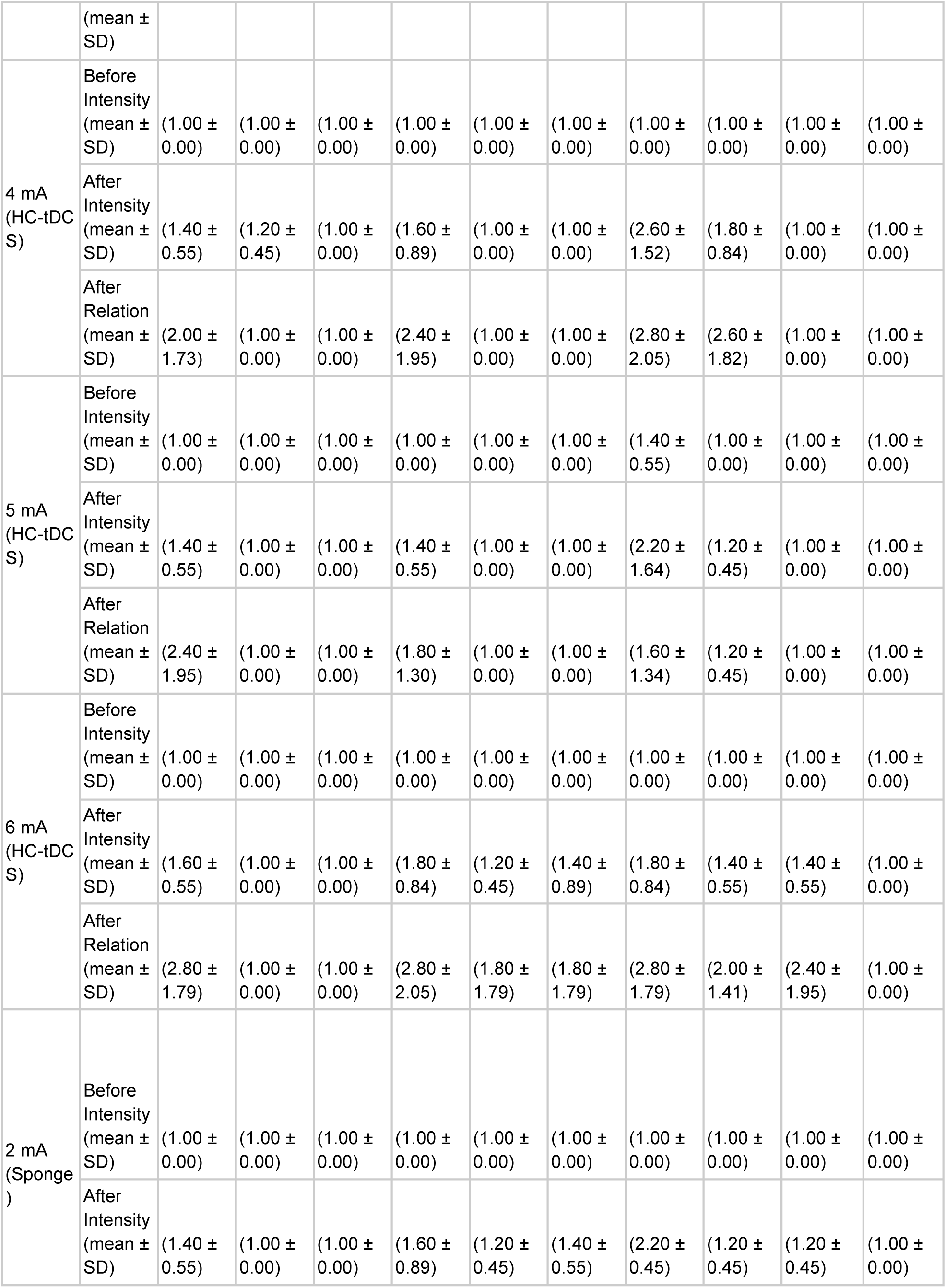
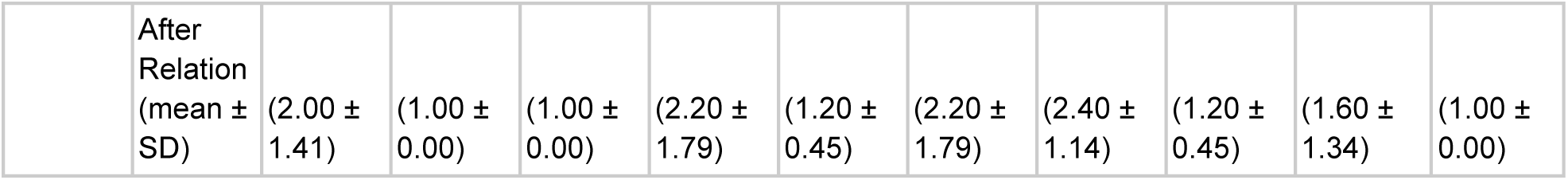

Analysis of Draize erythema scores had a significant main effect of current (β = 0.35, SE = 0.05, p < 0.0001), indicating that redness increased with higher stimulation currents. There was also a significant current-by-time interaction (β = –0.35, SE = 0.08, p < 0.0001). No significant main effects or interactions were found for the electrode side (all p > 0.4), and the three-way interaction was not significant (p = 0.58).

Skin temperature was measured at positions under the anode and cathode HC-tDCS electrodes, before and after stimulation for each target current (1-6 mA). There was no temperature increase of > .6° C in the electrode area relative to unstimulated forehead temperature for any condition tested across all subjects. No significant main effects of electrode side (β = 0.79°C, SE = 0.60, p = 0.19), time (before vs. after; β = 0.84°C, SE = 0.60, p = 0.17), or current (β = 0.19°C per unit mA, SE = 0.13, p = 0.16) was found on skin temperature. A significant interaction between the electrode side and time was observed (β = –1.73°C, SE = 0.85, p = 0.048). A significant three-way interaction among side, time, and current was also present (β = 0.65°C, SE = 0.26, p = 0.017).

### Human Trial: Distractor Task and Psychological Outcomes

*T*here was a significant main effect of block (β = 0.0035, SE = 0.00082, t(63,365) = 4.22, p < 0.001) and current-by-block interaction (β = –0.0015, SE = 0.00020, t(63,365) = –7.31, p < 0.001), but no main effect of current intensity(β = 0.0025, SE = 0.0019, t(63,365) = 1.36, p = 0.17), during HC-tDCS. Response times increased slightly over blocks, and the effect of current varied across time, with higher currents having a less pronounced increase. For accuracy of responses, no significant main effects of current (β = –0.0015, SE = 0.0010, t(382) = –1.39, p = 0.16) or block (β = –0.00089, SE = 0.00046, t(382) = –1.93, p = 0.054) were observed, and the current-by-block interaction was also not significant (β = 0.000070, SE = 0.00012, t(382) = 0.61, p = 0.54).

There were no significant differences or effects observed on any scored measure of acute mood (PANASPositive: current (β = 0.16, *p* = 0.631), time (β = –1.84, *p* = 0.323), interaction (β = –0.13, *p* = 0.790; PANASNegative: current (β = 0.05, *p* = 0.597), time (β = –0.46, *p* = 0.400),interaction (β = 0.07, *p* = 0.622); Anxiety: current (β = 0.03, *p* = 0.804), time (β = –0.59, *p* = 0.444), and interaction (β = –0.03, *p* = 0.866); Sadness: current (β = 0.05, *p* = 0.760), time (β = –0.54, *p* = 0.521), interaction (β = 0.20, *p* = 0.356); and Happiness: current (β = 0.26, *p* = 0.485), time (β = 0.11, *p* = 0.959), interaction (β = –0.57, *p* = 0.293))

## Discussion

Conventional tDCS at 2 mA produces peak brain electric fields of ≤1 V/m (36,37), with often still lower field strengths at the targeted brain regions (26,37,38). While animal and human intracranial studies demonstrated neural modulation at sub-V/m electric fields (9,40–43), higher electric fields enhance acute and plastic responses (11,12,44). Non-linear tDCS dose responses are special cases in TMS evoked responses (45, 46). Analysis of variance in clinical trial response supports higher electric fields increasing efficacy (14,15,47–49). Notwithstanding debate on optimal tDCS dosing, the exploration of the therapeutic window – increasing intensity while maintaining tolerability – is foundational. We show HC-tDCS even at 6 mA is as tolerated as 2 mA sponge-electrode tDCS while producing >3.5x higher electric fields. HC-tDCS supports a substantial expansion of the testable tDCS therapeutic window.

tDCS safety limits based on animal models of brain injury suggest thresholds of >60 mA (1). Given the tolerability of tDCS is associated with skin effects, technology to enhance dosing has focused on electrode design (17,18,21,22). HC-tDCS electrodes are novel in designing polarity-specific high-roughness redox layers (supporting faradaic charge transfer) and low-conductivity hydrogel (enhancing current uniformity and limiting diffusion from the redox layer), which are integrated into an comfortable articulated shape. The HC-tDCS stimulator is novel in a hybrid voltage-current controlled ramp up and low (7.5 V) compliance voltage. On the one hand, the HC-tDCS electrodes have low impedance minimizing the required compliance voltage. On the other hand the low voltage limits reactions at the redux electrode (≤1.5 for 1-6 mA, <30 min; Fig. 3a) and skin interfaces. The third HC-tDCS component is the protocol including preparing the skin with a designed wipe.

HC-tDCS use of adhesive hydrogel interface has associated advantages and limitations. The controlled adhesion supports skin-connection quality, and removes the need for headgear. Positioning is limited to below the hairline, though the bifrontotemporal montage itself enhances current delivery to the frontal cortex. Conventional bifrontal tDCS montages, such as F4-F4, produce diffuse current flow across the frontal cortex (39,48) as does bifrontotemporal HC-DCS (Fig. 4). Behavioural outcomes derive from network modulation (50–52) and/or functional targeting (53). The combination of high intensity and targeting can be achieved with HD-tDCS (54,55), but involves more complex equipment. HC-tDCS electrodes are by design single-use, preferable for sanitary applications.

## Notes

### Competing Interest Statement

Conflict of Interest
The City University of New York holds patents on brain stimulation with MB and NK as inventors. KD, MF, and NK consults for Ceragem Medical. MB has equity in Soterix Medical Inc. MB consults, provides expert witness support, received grants, assigned inventions, and/or served on the SAB of SafeToddles, Zabara Family Foundation, Boston Scientific, GlaxoSmithKline, Biovisics, Axonics, Mecta, Lumenis, Halo Neuroscience, Wave Neuroscience, Google-X, i-Lumen, Humm, Allergan(Abbvie), Apple, Ybrain, Ceragem, Ceragem Clinical, Remz. MB is supported by grants from Harold Shames and the National Institutes of Health: NIH-NIDA UG3DA048502, NIH-NIGMS T34 GM137858, NIH-NINDS R01 NS112996, NIH-NINDS R01 NS101362, and NIH-G-RISE T32GM136499.

### Funding

- NIH: UG3DA048502,, T34 GM137858, R01 NS112996, R01 NS101362, T32GM136499

